# A Role for Steroid 5 alpha-reductase 1 in Vascular Remodelling During Endometrial Decidualisation

**DOI:** 10.1101/2022.05.30.493728

**Authors:** I.W. Shaw, P.M. Kirkwood, D. Rebourcet, F.L. Cousins, R.J. Ainslie, D.E.W. Livingstone, L.B. Smith, P.T.K. Saunders, D.A. Gibson

**Author notes:** to whom correspondence should be addressed., Centre for Inflammation Research, The University of Edinburgh, Institute for Regeneration and Repair, Edinburgh Bioquarter, Edinburgh.

## Abstract

Decidualisation is the hormone-dependent process of endometrial remodelling that is essential for fertility and reproductive health. It is characterised by dynamic changes in the endometrial stromal compartment including differentiation of fibroblasts, immune cell trafficking and vascular remodelling. Deficits in decidualisation are implicated in disorders of pregnancy such as implantation failure, intra-uterine growth restriction, and pre-eclampsia.

Androgens are key regulators of decidualisation that promote optimal differentiation of stromal fibroblasts and activation of downstream signalling pathways required for endometrial remodelling. We have shown that androgen biosynthesis, via 5α-reductase-dependent production of dihydrotestosterone, is required for optimal decidualisation of human stromal fibroblasts *in vitro*, but whether this is required for decidualisation *in vivo* has not been tested.

In the current study we used steroid 5α-reductase type 1 (SRD5A1) deficient mice (*Srd5a1-/-* mice) and a validated model of induced decidualisation to investigate the role of SRD5A1 and intracrine androgen signalling in endometrial decidualisation. We measured decidualisation response (weight/proportion), transcriptomic changes, and morphological and functional parameters of vascular development. These investigations revealed a striking effect of 5α-reductase deficiency on the decidualisation response. Furthermore, vessel permeability and transcriptional regulation of angiogenesis signalling pathways, particularly those that involved vascular endothelial growth factor (VEGF), were disrupted in the absence of 5α-reductase. In *Srd5a1-/-* mice, injection of dihydrotestosterone co-incident with decidualisation restored decidualisation responses, vessel permeability, and expression of angiogenesis genes to wild type levels.

Androgen availability declines with age which may contribute to age-related risk of pregnancy disorders. These findings show that intracrine androgen signalling is required for optimal decidualisation in vivo and confirm a major role for androgens in the development of the vasculature during decidualisation through regulation of the VEGF pathway. These findings highlight new opportunities for improving age-related deficits in fertility and pregnancy health by targeting androgen-dependent signalling in the endometrium.

## Introduction

Decidualisation is a fundamental step in the establishment of pregnancy that involves coordinated remodelling of the endometrial stroma. It is a hormone-dependent process characterised by differentiation of endometrial stromal fibroblasts (hESF) (Dunn et al., 2003), immune cell trafficking (Yang et al., 2019) and vascular remodelling (Plaisier, 2011). Deficits in decidualisation are implicated in disorders of pregnancy such as implantation failure, intrauterine growth restriction, and pre-eclampsia (Dunk et al., 2008, Gibson et al., 2016b). Risk of pregnancy disorders increases with age which may be associated with age-related decline in hormone production or availability.

Androgens are key regulators of decidualisation that promote optimal differentiation of stromal fibroblasts and activation of downstream signalling pathways required for endometrial remodelling. Androgen supplementation enhances secretion of decidualization markers in hESF *in vitro* (Narukawa et al., 1994). Exogenous dihydrotestosterone (DHT) enhances and maintains decidualisation responses in mice, an effect which is attenuated by co-administration of the AR antagonist flutamide (Zhang and Croy, 1996). Endometrial fibroblasts in both mouse and human endometrium express androgen receptor (AR) and their function is altered by AR-dependent signalling (Marshall et al., 2011, Simitsidellis et al., 2019, Simitsidellis et al., 2016). Using assays that combined *in vitro* decidualisation and knockdown of receptors Cloke et al. reported that AR regulated distinct decidual gene networks involved in cytoskeletal organization, cell motility and regulation of the cell cycle (Cloke et al., 2008). Using a pharmacologic approach, we found that flutamide attenuated the expression of decidualisation and endometrial receptivity markers in hESF (Gibson et al., 2016a). Together, these studies confirm the importance of androgens and AR-dependent signalling for optimal decidualisation responses.

In women, androgens and their precursors are abundant in the circulation but production declines with age (Zumoff et al., 1995, Labrie et al., 1997). Circulating concentrations of the most potent androgen dihydrotestosterone (DHT) are low but this is because DHT is primarily a product of local metabolism within target tissues in women; mediating its affects via intracrine signalling (Rothman et al., 2011, Labrie et al., 2017). We have shown that DHT is actively produced by endometrial stromal fibroblasts during decidualisation via expression of the enzyme SRD5A1 and that the abundance of DHT is affected by availability of steroid precursors such as dehydroepiandrosterone (DHEA) (Gibson et al., 2018a, Gibson et al., 2016a, Gibson et al., 2018b, Gibson et al., 2016b). When we increased intracrine DHT signalling by supplementing with DHEA we found that decidualisation responses were enhanced and expression of implantation markers were also increased. Androgen precursor availability and intracrine androgen signalling can therefore dictate the extent of decidualisation responses.

To date, a role for intracrine androgen signalling in regulating decidualisation *in vivo* has not been rigorously investigated. Previous studies have been hampered by the developmental uterine defects reported in global AR knock-out female mice (Walters et al., 2016). In the current study, we have used mice which are homozygous for a recessive knock-out mutation in the gene encoding the type 1 steroid 5α-reductase (SRD5A1) enzyme (*Srd5a1*^*-/-*^) which lack the capacity to convert T to DHT (Mahendroo et al., 1996). Previously, Srd5a1^-/-^ mice have been shown to have reduced fertility due to oestrogen-induced foetal death and a parturition defect caused by lack of 5α-androstan-3α,17β-diol, another product of SRD5A1 (Mahendroo et al., 1997). The potential role of this enzyme and the contribution of 5α-reduced androgens to decidualisation has not been tested, but it is plausible that the reduced fecundity reported in Srd5a1^-/-^ mice (Mahendroo and Russell, 1999) may in part be due to decidualisation defects as a result of intracrine androgen deficiency. To investigate this, we have assessed the impact of 5α-reductase deletion or inhibition in a mouse model of induced decidualisation.

We show that 5α-reductase deficiency leads to impaired decidualisation, structural and functional changes to decidual blood vessels, and transcriptomic changes affecting angiogenesis signalling pathways. Deficits in decidualisation in Srd5a1^-/-^ mice were reversed by restoration of androgen signalling through exogenous DHT administration. We conclude that intracrine androgens are necessary for optimal decidualisation and vascular remodelling required for establishment of pregnancy.

## Materials and Methods

### Mice

All mouse work was performed in accordance with the Animals (Scientific Procedures) Act 1986 under UK law and were bred on a C57Bl/6J or C57Bl6/JCrl background. The Srd5a1^-/-^ mouse line was obtained from the Jackson Laboratories (https://www.jax.org/strain/002793). The original paper describing the generation of mice with a null allele was published by Mahendroo, Cala, and Russel (Mahendroo et al., 1996). Due to parturition defects in homozygous mice heterozygote crosses were used to maintain the colony and produce *Srd5a1*^*-/ -*^ progeny. Wildtype (WT) mice were either littermates or purchased from Charles River Laboratories (Tranent, Scotland). Expression of 5α-Reductase isozymes (types 1-3) was assessed in uterine tissues by qPCR (Supplementary Figures 1). *Srd5a1* was absent in uterine tissues from knockout mice, and consistent with previous reports (Mahendroo et al., 1996), *Srd5a2* was not detected in either wildtype or knockout mice. While *Srd5a3* was detected its expression was decreased with decidualisation (Supplementary Figures 1).

**Figure 1.**
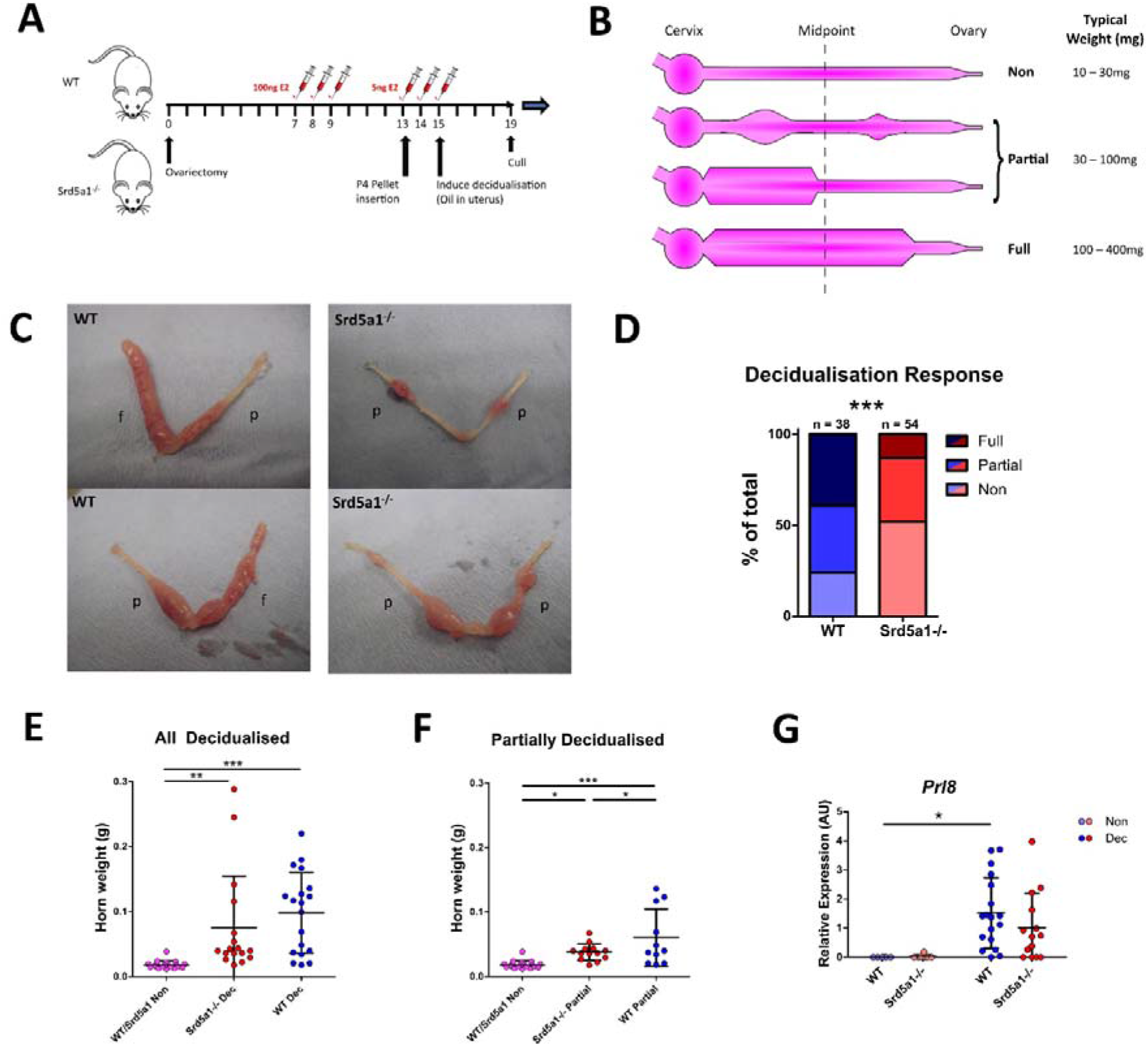
Decidualisation response is impaired in Srd5a1^-/-^ mice. A) Schema for the decidualisation induction model. E2 = oestradiol. P4 = progesterone. B) Graphic depicting a uterine horn in various stages of decidualisation. The horn is thicker where a decidualisation reaction has occurred. A horn is deemed fully decidualised if an uninterrupted stretch of decidualisation has occurred over at least half of the horn length. Partial decidualisation is any decidualisation response less than this, often occurring as disparate nodes along the horn. C) Representative images of uterine horns from wild type (WT) and Srd5a1^-/-^ animals, with the classification of each horn given; p = partial decidualisation, f = fully decidualised. D) Contingency table depicting proportion of horn responses in each genotype. E,F) Wet weight of uterine horns in WT and Srd5a1^-/-^ mice at time of tissue recovery (4 days following decidualisation induction). Non-decidualised horns have been pooled. E compares all horns with a decidualisation reaction (Full & Partial), whereas F shows analysis of those with a partial reaction. G) Expression of *Prl8* via qPCR in uterine horns of WT and Srd5a1^-^/^-^ mice following decidualisation induction. Plots in D analysed by chi-squared test for trend; E, F analysed by one-way ANOVA with Newman-Keuls *post-hoc* tests; and G analysed by two-way ANOVA with Bonferroni post-hoc tests; *** p < 0.001; ** p < 0.01; * p < 0.05.

### Mouse model of induced decidualisation

Mice were ovariectomised to remove endogenous ovarian steroids at day 0 (d0), then administered oestradiol (E2, Sigma-Aldrich, Poole) by subcutaneous injection daily on d7-9 (5ug.kg^-1^ in 200ul sesame oil) and on d13-15 (0.25ug.kg^-1^ in 200ul sesame oil) with a P4 pellet inserted on d13. On day 15 a decidualisation stimulus was administered by intrauterine injection of 20ul sesame oil into the uterus via transvaginal delivery using a non-surgical embryo transfer device (NSET). Animals were sacrificed 4 days later and uterine tissues collected (Fig. 1A, (Cousins et al., 2014)). The decidualisation response was immediately scored in each horn as outlined in Fig. 1B. Specifically a non-decidualised horn had no decidualisation reaction whatsoever (Non), whereas a fully decidualised horn had continuous decidualisation along the majority (>50%) of its length (Full). Partial decidualisation is any level of decidualisation between these two extremes (Fig. 1B). Collectively, samples designated Full or Partial are referred to as decidualised (Dec).

Finasteride (Sigma-Aldrich, Poole) was dissolved in 100% EtOH, then diluted to 5% v/v in sesame oil to a final concentration of 12.5mg.mL^-1^ and 100uL (50 mg.kg^-1^) was administered via intraperitoneal injection daily from the point of the decidualisation stimulus onwards, i.e. d15-18, as show in Figure S1A. Finasteride is a synthetic 4-azasteroid that competitively inhibits both type I and II 5α-reductase enzymes in rodents (Livingstone et al., 2015, Thigpen and Russell, 1992).

DHT (Sigma-Aldrich, Poole) was dissolved in 100% ethanol (EtOH) then diluted to 5% v/v in 0.4% w/v methylcellulose solution for a final DHT concentration of 2mg.mL^-1^. 100uL (8mg.kg^-1^) was injected subcutaneously on d15 at the point of decidualisation stimulus. At the end of experiments mice were terminated by exposure to rising concentrations of CO_2_ combined with cervical dislocation.

### Histology and immunolabelling

Uterine tissue for histology was fixed in 4% paraformaldehyde then dehydrated and embedded in paraffin blocks and sectioned according to standard protocols. Protocols for 3,3’-diaminobenzidine (DAB) and immunofluorescence staining were performed as previously reported (Kirkwood et al., 2021, Simitsidellis et al., 2016). Briefly, slides were dewaxed, rehydrated, and subjected to antigen retrieval in pH 6.0 citrate buffer using a microwave. Following a wash in phosphate buffered saline (PBS) slides were incubated with 3% H_2_O_2_ solution in PBS for 15min to remove endogenous peroxidase activity. Following several PBS washes slides were incubated with normal goat serum solution (NGS, PBS containing 20% goat serum and 0.05% (w/v) bovine serum albumin) to block non-specific interactions. Anti-CD31 primary antibody (Abcam, ab28364, 1:500 dilution) in NGS was added and incubated overnight at 4°C. Following several washes, slides were incubated with goat-raised peroxidase-conjugated secondary antibody (Abcam, ab7171, 1:500 dilution) for 1hr at room temperature. Following several washes, slides were incubated with either DAB reagent or Opal red 570 reagent for 10 minutes or until sufficiently developed. After a brief wash, slides were counterstained either with haematoxylin and eosin as per standard procedures, or with DAPI before being mounted with coverslips.

Slides were imaged using a Zeiss Z1 microscope, or a Zeiss AxioScan Slide Scanner using conventional fluorescent setups.

### Vessel quantification

To quantify vessels, regions of non-decidualised endometrium (Non) or decidua (Dec samples) were selected digitally on 20X magnification slide scans of CD31-labelled uterus sections and overlaid with a grid with edge length 250 pixels (∼ 28μm). The intersections on the grid were sequentially classified as either vessel or not vessel, and the total proportion of vessel to non-vessel intersections was used as a measure of vessel count. Additionally, each time a vessel was identified, it was given an irregularity score equalling the number of intersections on a tree used to trace out the vessel’s cross-sectional shape. For example, a circular or or ‘L’ shaped cross-section would have a score of 0, a ‘T’ shape a score of 1, an ‘H’ shape a score of 2, etc. Irregularity scores were averaged across all observations.

### RNA extraction

Uterine samples for RNA analysis were collected in RNALater solution (Qiagen, Hilden, Germany) and stored at -80°C until tissue extraction. For decidualised samples, only regions of uterus containing decidualised endometrium were selected. Samples were thawed and homogenised in 1 mL Trizol reagent using a TissueLyser machine (6min, 25Hz). Homogenate was transferred to MaXtract High Density tubes along with 200ul chloroform and 100ul RNase free water, shaken vigorously and centrifuged. The aqueous layer was added to RNeasy spin columns (Qiagen, Hilden, Germany) and processed as per manufacturer’s instructions.

### Quantitative RT-PCR

cDNA was synthesised from 100ng.uL^-1^ RNA using Superscript VILO cDNA synthesis kit (Thermo Fisher Scientific Life Sciences, Schwerte, Germany) as per the manufacturer’s instructions. Thermal cycler settings were 25°C for 10min, 42°C for 60min, 85°C for 5min. Primers were designed using Universal Probe Library Assay Design Center (Roche Applied Science, Burgess Hill, UK) or, when this was discontinued, using Neoformit (https://primers.neoformit.com/ Accessed: 6/4/22; sequences in Supplementary Data 1). Primers were synthesised by Eurofins MWG Operon (Ebersberg, Gemany). Reactions were run in duplicate on a Quantstudio 5 384-well PCR machine (Thermo Fisher Scientific Life Sciences, Schwerte, Germany) using the following settings: 95°C for 10min then 40 cycles of 95°C for 15s and 60°C for 1min. A serial dilution of a standard cDNA solution (consisting of a mix of all samples tested) was used to plot a standard curve and expression levels interpolated from this. Data was processed using QuantStudio Design and Analysis Software.

### Nanostring gene expression analysis and data processing

Samples were analysed on the NanoString nCounter Analysis System using a NanoString Mouse Pancancer Pathways Panel kit (Nanostring, Edinburgh, UK). RNA at 20ng.uL^-1^ was used. Processing was performed by the Host and Tumour Profiling Unit Microarray Services, Institute of Genetics and Cancer, University of Edinburgh, as per the manufacturer’s instructions. Briefly, the reporter codeset was mixed with hybridisation buffer and added to each of the RNA samples, followed by the capture codeset. This mix was hybridised at 65°C for 18hr. Following hybridisation samples were loaded into the provided cartridges on the nCounter prep station and processed using the High Sensitivity protocol. Cartridges were then sealed and read using the digital analyser on the max setting. No QC flags were registered for any of the samples.

Data was processed using the nSolver analysis software. Data was normalised using a combination of positive controls and housekeeping genes. Firstly, background was subtracted using a background value of the mean plus two standard deviations of the negative control values. Normalisation using positive controls and housekeeper genes was performed using the geometric mean to compute the normalisation factor, and the manufacturer-provided list of housekeeper genes was used excluding the following genes based on the results of the geNorm algorithm in the Nanostring Advanced Analysis modules: *Hprt, Alas1, G6pdx, Gusb, Ppia*.

Normalised counts data was then analysed for differential gene expression using R, specifically the edgeR analysis workflow. Dispersions were estimated with the *estimateDisp()* command and fit to the design model using *glmQLFit()*, before testing for differential expression using *glmQLFTest()*. Graphs were produced using the ggplot2 package. Finally, to perform gene ontogeny (GO) analysis we utilised the clusterProfiler analysis workflow. Ensembl IDs were mapped from the Bioconductor org.Mm.eg.db genome annotation and enrichment assessed using the *enrichGO()* command with the ‘universe’ set as the genes included in the Nanostring Pancancer Pathways panel. Graphs were produced using the *dotplot ()* and *cnetplot ()* commands from the enrichplot R package.

### Comparison of Nanostring gene expression with scRNAseq gene expression

Normalised counts and differential gene expression data was further analysed in the context of publicly available single cell RNA sequencing (scRNAseq) data, accessed from NCBI’s Gene Expression Omnibus (Edgar et al., 2002) through GEO Series accession numbers GSE160772 and GSE198556. These scRNAseq data defined the transcriptomic profile of over 20,000 perivascular, fibroblast and epithelial cell types present in the mouse uterus. The AverageExpression() command (Seurat) was used to calculate the mean expression of Nanostring DEgenes in perivascular, fibroblast and epithelial cell clusters from scRNAseq data. Gene expression matrices were normalised and clustered heatmaps produced using the pheatmap package. Expression values shown are scaled but not centred. This comparison allowed us to attribute the expression of genes that were found to be differentially expressed between Srd5a1^-/-^ and WT genotypes to certain cell fractions in the mouse endometrium.

### Creation and imaging of resin casts

Resin casts were prepared using a method previously validated in male mice (Rebourcet et al., 2016). Briefly, female mice (Non and Dec) were culled using a terminal dose of sodium pentobarbital (150 mg/kg, ip) and perfusion fixation of the vasculature was achieved via the left ventricle. Heparinized PBS (heparin, 20 U/mL) was infused at 6 mL/min for 2 minutes. Low-viscosity resin (10 mL; Microfil MV-122; Flow Tech Inc) was prepared according to the manufacturer’s instructions and then infused via the left ventricle. Uterine tissues were recovered, trimmed, fixed in paraformaldehyde and embedded in 1.5% low melting point agarose (Invitrogen, UK), dehydrated in methanol (100%; 24 hours) and optically cleared in benzyl alcohol:benzyl benzoate (1:2 v/v; 24 hours). Imaging was carried out as described in Rebourcet et al. with a positive control tissue being a previously prepared mouse testis (Rebourcet et al., 2016).

### Evans Blue assay of vascular permeability

Evans Blue (EB) dye binds to albumin in the blood so its presence in tissues, which can be quantified spectrophotometrically, implies a breakdown in the endothelial barrier. The Evans Blue dye (Sigma-Aldrich, Poole) was dissolved in sterile saline solution to a concentration of 0.5% w/v, and 200uL was injected i.v. (via tail vein) 30-90 minutes prior to cull via cervical dislocation. The interval between EB injection and cull was recorded and found not to associate with dye retention in tissues.

To quantify the amount of Evans blue in the tissues, following sacrifice a portion of the uterus was taken, precisely weighed, and transferred to 500μl of formamide. The gross appearance of the tissue was recorded using a digital camera. For decidualised samples, only sections of uterus containing decidualised endometrium were selected. Samples were incubated for 24h at 55°C to extract the dye, then tissue pieces were discarded, the solution centrifuged to remove debris, and absorbance read in triplicate at 610nm.

### Data processing and statistical analysis

Unless otherwise stated, data was processed using Microsoft Excel and Python. Statistics were performed using either Python or Graphpad Prism 5.

In the analysis of Nanostring data, two-way ANOVAs were performed on all genes found to be expressed and ANOVA results corrected for multiple comparisons using the Benjamini and Hochberg method (Benjamini and Hochberg, 1995). The false discovery rate (FDR) was 0.05.

In the analysis of Evans Blue absorbance data, absorbance readings are shown as a ratio relative to the mean value of WT or vehicle-treated WT data. In the case of data from Figure 2, ratios have been calculated separately for partially and fully decidualised horns and the results combined for the subsequent statistical analysis.

**Figure 2.**
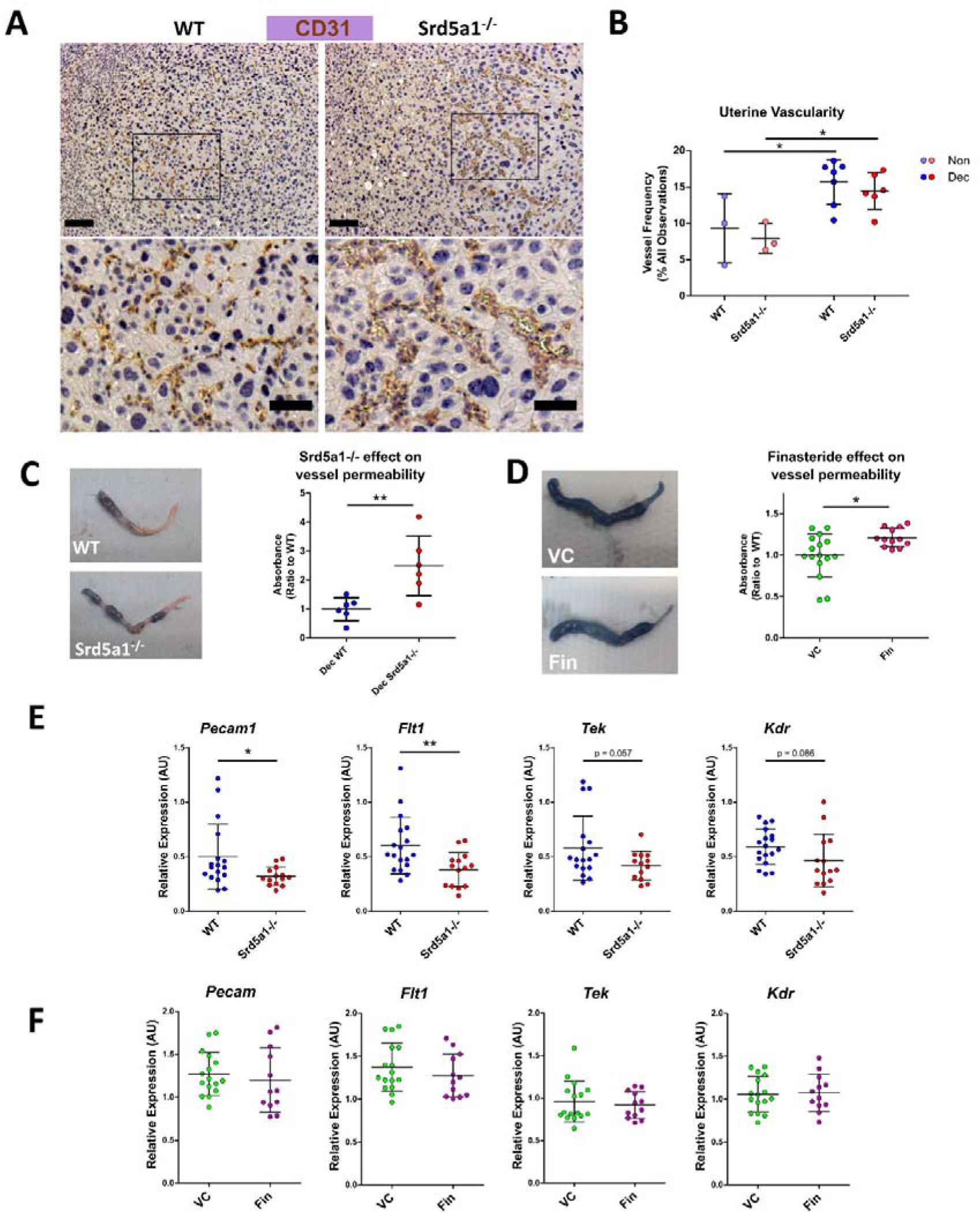
Evidence of a vascular phenotype in Srd5a1^-/-^ mice following induction of decidualisation. A) Representative images of CD31^+^ cells in WT and Srd5a1^-/-^ decidua. Zoomed-in views are shown from regions indicated by boxes. Scale bars are 100μm and 50μm respectively. B) Quantification of vessel density in in WT and Srd5a1^-/-^ non-decidualised endometrium and decidua. C, D) Images of decidualised uterine horns following intravenous injection of Evans Blue and respective quantification of absorbance in uterine tissue extracts in decidualised horns of WT and Srd5a1^-/-^ (C) or vehicle control (VC) and finasteride (Fin) treated mice (D). E,F). Expression of angiogenic genes via qPCR in decidualised tissue (partial and full) from WT and Srd5a1^-^/-mice (E), or VC and Fin mice (F). Plots in B analysed by two-way ANOVA with Bonferroni *post-hoc tests;* C,D,E,F analysed by two-tailed t test. ** p < 0.01; * p < 0.05.

Statistical tests performed for other comparisons are stated in figure legends and the text.

## Results

### 5*α*-Reductase deficiency impairs decidualisation in mouse uterus

To investigate the functional requirement for 5α-reductase during decidualisation we induced decidualisation in WT and Srd5a1^-/-^ mice using a previously validated protocol (Fig. 1A; (Cousins et al., 2014)). We used a ternary system to classify the extent of decidualisation. Uterine horns were classified as non-, partially- or fully-decidualised (Non, Partial, Full, respectively; Fig. 1B). For the purposes of analysis individual uterine horns were treated as separate biological entities. In instances where only the decidualised regions of the uterus are analysed Partial and Full horns were grouped together: referred to as ‘decidualised’ (Dec). Decidualisation was associated with a significant increase in *Srd5a1* mRNA expression in wildtype mice which was absent in knockout mice (Supplementary Figures 1). Decidualisation was observed in both genotypes, however Srd5a1^-/-^ uteri had a noticeably impaired decidualisation response compared with those of WT. Specifically uteri from Srd5a1^-/-^ mice had fewer ‘Full’ horns and decidualised areas in ‘Partial’ horns were smaller (Fig 1C). When quantified and compared statistically we were able to reject the hypothesis that the number in each of the decidualisation response groups is the same between genotypes, which supports our interpretation that decidualisation is impaired in Srd5a1^-/-^ mice (Fig 1D).

In line with expectations uterine weight increased significantly in Dec horns compared to non-decidualised horns in both WT and Srd5a1^-/-^ mice (Fig 1E). Uterine horn weights of partially decidualised horns from WT mice were significantly heavier than those from Srd5a1^-/-^ females (Fig 1F). The expression of mRNAs encoded by *Prl8*, a marker of decidualisation, was significantly increased in Dec compared to Non horns in WT mice. A similar, non-significant, trend for increased *Prl8* mRNA expression was observed in Srd5a1^-/-^ mice (Fig 1G).

To complement these findings we tested the impact of transiently blocking 5α-reductase activity in WT mice using the pharmacologic 5α-reductase inhibitor finasteride (for 4 days from days 15-18; Supplementary Figures 2A). Notably this treatment regime did not significantly affect the decidualisation response, uterine horn weights, nor expression of *Prl8* (Supplementary Figures 2B,C,D). Thus transient inhibition of 5α-reductase under this paradigm was insufficient to replicate the effects of the global *Srd5a1* knock-out on decidualisation responses.

### 5*α*-reductase is required for normal endothelial barrier function and vascular remodelling during decidualisation

To investigate if impaired decidualisation in Srd5a1^-/-^ mice was also associated with deficient vascular morphology and function we stained blood vessels with the endothelial cell marker CD31. Observational analysis of tissue sections suggested blood vessels in mutant mice appeared less numerous and more dilated compared to WT (Fig 2A). To quantify the vessels, counts were performed on samples fluorescently labelled for CD31 (Supplementary Figures 3A). This analysis revealed a significant increase in vessel frequency in decidualised compared to non-decidualised uterine horns (Fig 2B). The mean vessel frequency in Srd5a1^-/-^ decidualised uteri was lower than in WT but no statistically significant difference was detected (Fig 2B). Vessel irregularity was also quantified, revealing a significant increase with decidualisation, but no distinct effect of genotype (Supplementary Figures 3B).

**Figure 3.**
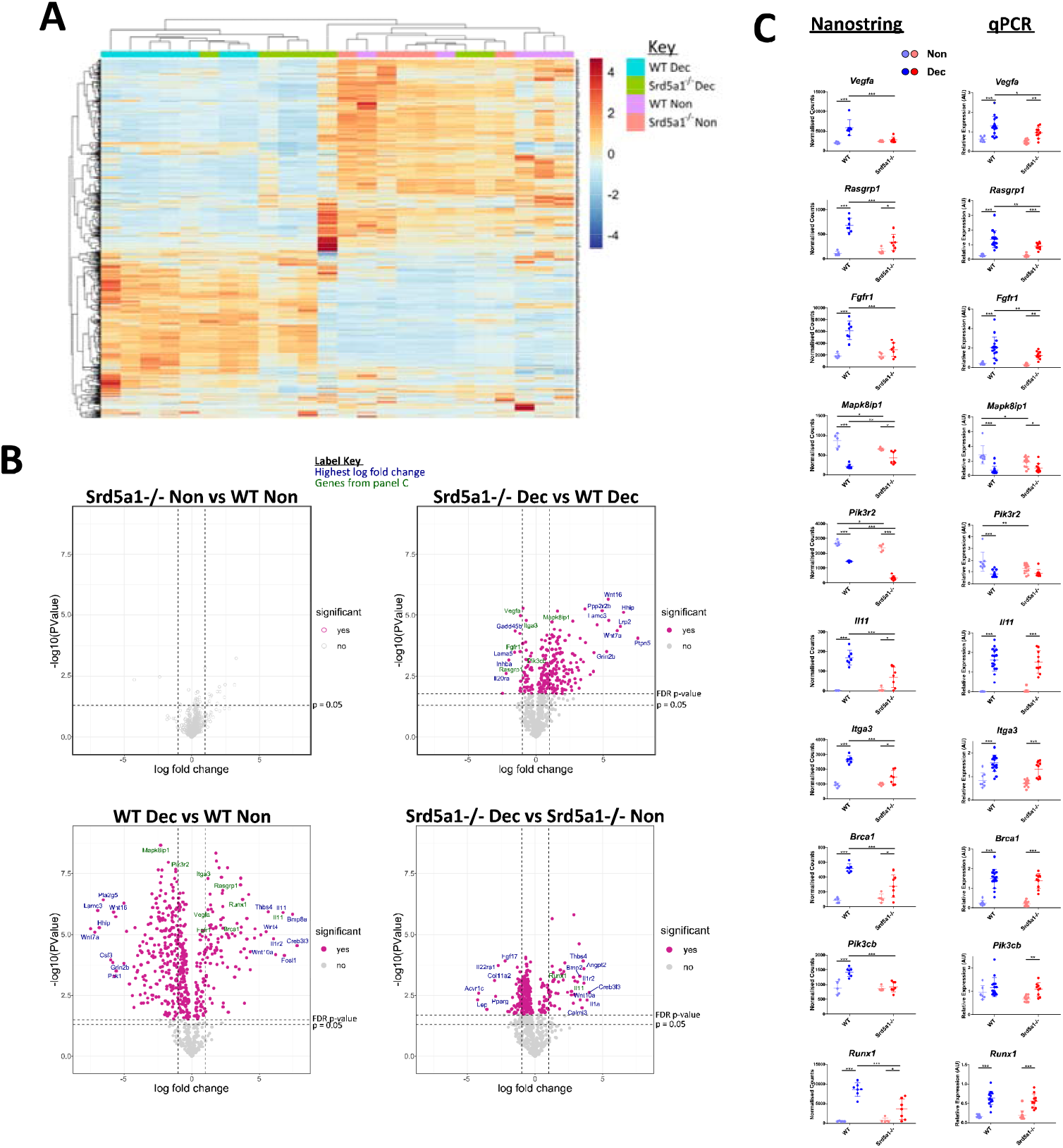
Effect of 5α-reductase deficiency on global gene expression during decidualisation. A) Unsupervised clustering and heatmap of WT and Srd5a1^-/-^ decidualised and non-decidualised tissue RNA analysed with Nanostring Pancancer Pathways gene expression panel. The major gene expression differences occur with decidualisation. B) Volcano plots for the pairwise comparisons between experimental groups, as indicated. Significantly differentially expressed (DE) genes at a false discovery rate (FDR) of 0.05 are coloured pink. C) Selection of genes changing significantly with either genotype, or interaction of genotype and decidualisation, by two-way ANOVA (FDR = 0.05). Count values in the Nanostring data and respective validation by qPCR in groups with additional samples is shown. Plots in C analysed by two-way ANOVA with Bonferroni *post-hoc* tests. * p < 0.05; ** p < 0.01; *** p < 0.001.

We attempted to map the three-dimensional architecture of vessels using the resin cast method, however accurate casts could not be obtained which we believe is due to the high permeability of vessels in the decidualised tissues (Supplementary Figures 4). This finding and observations from histological slides of dilated vessels in the decidual tissue of Srd5a1^-/-^ mice led us to consider whether vessel permeability might be altered as a consequence of ablation of the gene in mice.

**Figure 4.**
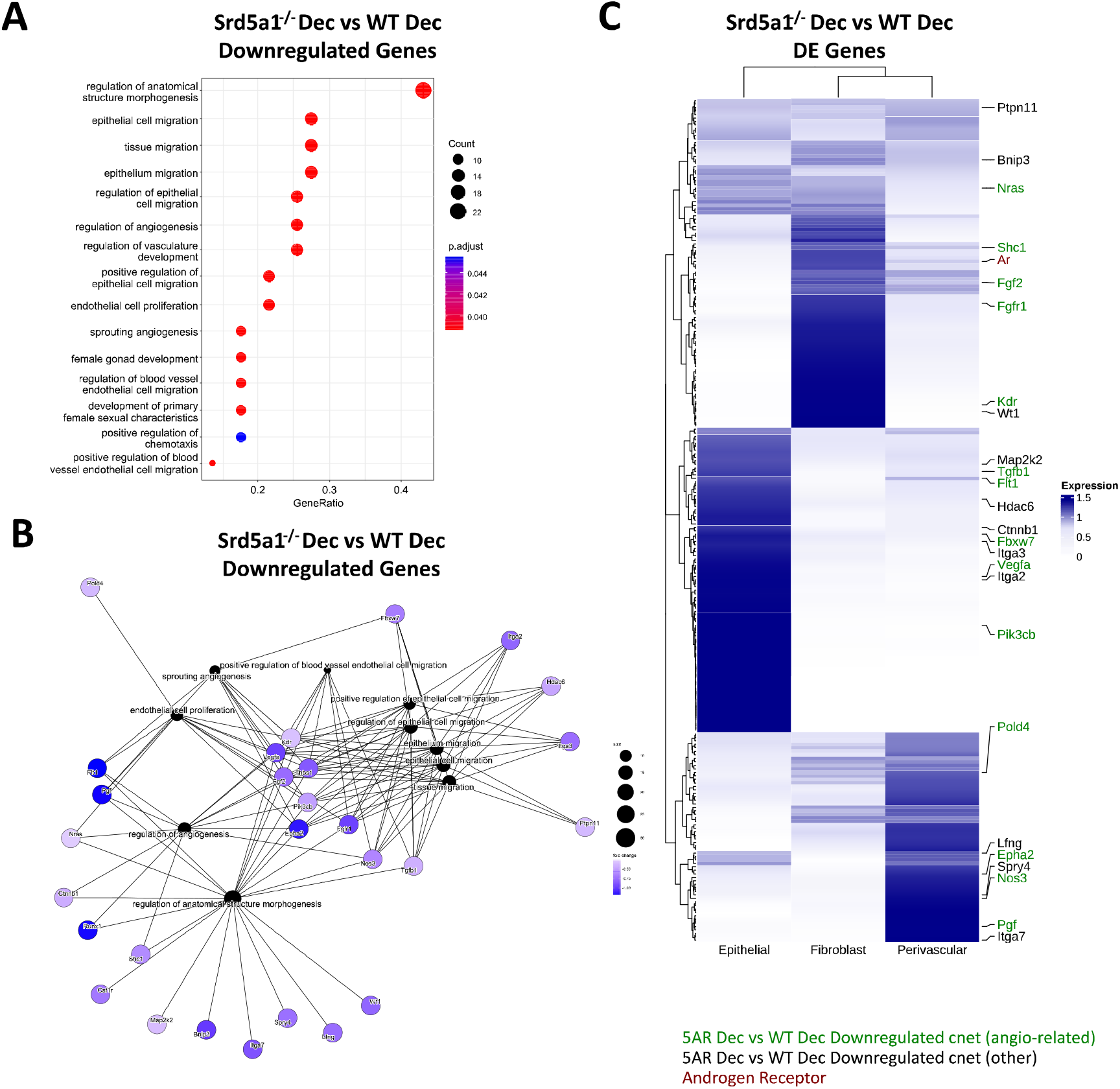
Gene networks associated with defective decidualisation in Srd5a1^-/-^ mice. A) Biological process gene ontogeny (GO) of genes significantly downregulated in Srd5a1^-/-^ vs WT decidualised uterine horns. All significant terms shown. B) Category netplot (cnet) embedding indicating genes shared between GO terms for genes significantly downregulated in Srd5a1^-/-^ vs WT decidualised uterine horns. C) Heatmap showing expression of top DE genes from the Srd5a1^-/-^ compared to WT decidualised uterine horns and their relative abundance in uterine cell subsets; epithelial, fibroblast and perivascular, as derived from annotation in previously published scRNAseq dataset of mouse uteri. Genes appearing in the cnet plot in B are labelled and colour coded based on whether they link to angiogenesis-related GO terms (green) or not (black).

To quantify whether these morphological changes lead to functional deficiency in endothelial barrier integrity, we investigated vascular permeability using Evans Blue assay. We observed a significant increase in permeability of decidualised uterine horns from Srd5a1^-/-^ compared to WT mice (Fig 2C). In mice treated with finasteride, similar trends were observed, and there was a significant increase in permeability in decidualised horns following finasteride treatment (Fig. 2D).

### Changes in angiogenic gene expression during decidualisation are attenuated in Srd5a1^-/-^ mice

To investigate the potential molecular basis for the vascular phenotype observed in Srd5a1^-/-^ uteri, we quantified the expression of key angiogenic genes in decidualised horns from WT and Srd5a1^-/-^ mice. A significant decrease in the expression of *Pecam1* and *Flt1* was detected in Srd5a1^-/-^ uteri compared to WT (Fig 2E). The mean expression of *Tek* (TIE2/Angiopoietin-1 receptor) and *Kdr* (VEGFR2) was also decreased but this was not statistically significant (Fig 2E). When gene expression was analysed in WT mice treated with finasteride there was no detectable change in the expression of the same panel of angiogenic genes (Fig. 2F).

### 5α-Reductase deficiency attenuates decidualisation-induced changes in gene expression

To discover genes and signalling pathways that are altered by deletion of *Srd5a1* in decidualised endometrium we analysed the transcriptome of Non and Dec uterine tissue from WT and Srd5a1^-/-^ mice using the Nanostring Pancancer Pathways panel. To visualise the general patterns of gene expression, count values were subjected to unsupervised hierarchical clustering and visualised on a heat map (Fig. 3A). Clustering of samples forms two major arms, one containing exclusively Dec samples and the other mostly Non. These findings are consistent with decidualisation being a major driver of changes in gene expression in these samples. Notably, within the Dec arm Srd5a1^-/-^ samples largely clustered together and closer to the Non arm, and several decidualised Srd5a1^-/-^ samples are found within the Non arm, suggesting a transcriptional phenotype in Srd5a1^-/-^ Dec uterus which was not the same as in the WT Dec tissue.

To interrogate the differential expression of genes between the two genotypes, we performed four ratio comparisons: Srd5a1^-/-^ Non vs WT Non; Srd5a1^-/-^ Dec vs WT Dec; WT Dec vs WT Non; and Srd5a1^-/-^ Dec vs Srd5a1^-/-^ Non (Fig. 3B). This analysis revealed no significantly differentially expressed (DE) genes between genotypes in non-decidualised tissue (Srd5a1^-/-^ Non vs WT Non), suggesting a minimal role for 5α-reductase in regulating uterine gene expression in the absence of decidualisation. In stark contrast, decidualisation in WT mice induced significant changes in the expression of many genes: 507 or 66% of the total analysed (WT Dec vs WT Non; Table 1). Interestingly, the majority (71%) of the DE genes in this panel were down-regulated by decidualisation. In comparison with the WT data, gene expression changes in Srd5a1^-/-^ decidualised tissues (Srd5a1^-/-^ Dec vs Srd5a1^-/-^ Non) were greatly suppressed, with fewer DE genes (299, 39%) and smaller changes in expression level for both up- and down-regulated genes (Table 1). When the expression of genes in decidualised uteri of WT and Srd5a1^-/-^ mice was directly compared (Srd5a1^-/-^ Dec vs WT Dec), many significantly DE genes are observed (254, 33%), most of which are up-regulated in Srd5a1^-/-^ compared to WT (80%). Notably when significantly DE genes were considered 95% of the up-regulated genes in this comparison are downregulated in the WT Dec vs WT Non comparison.

**Table 1:**
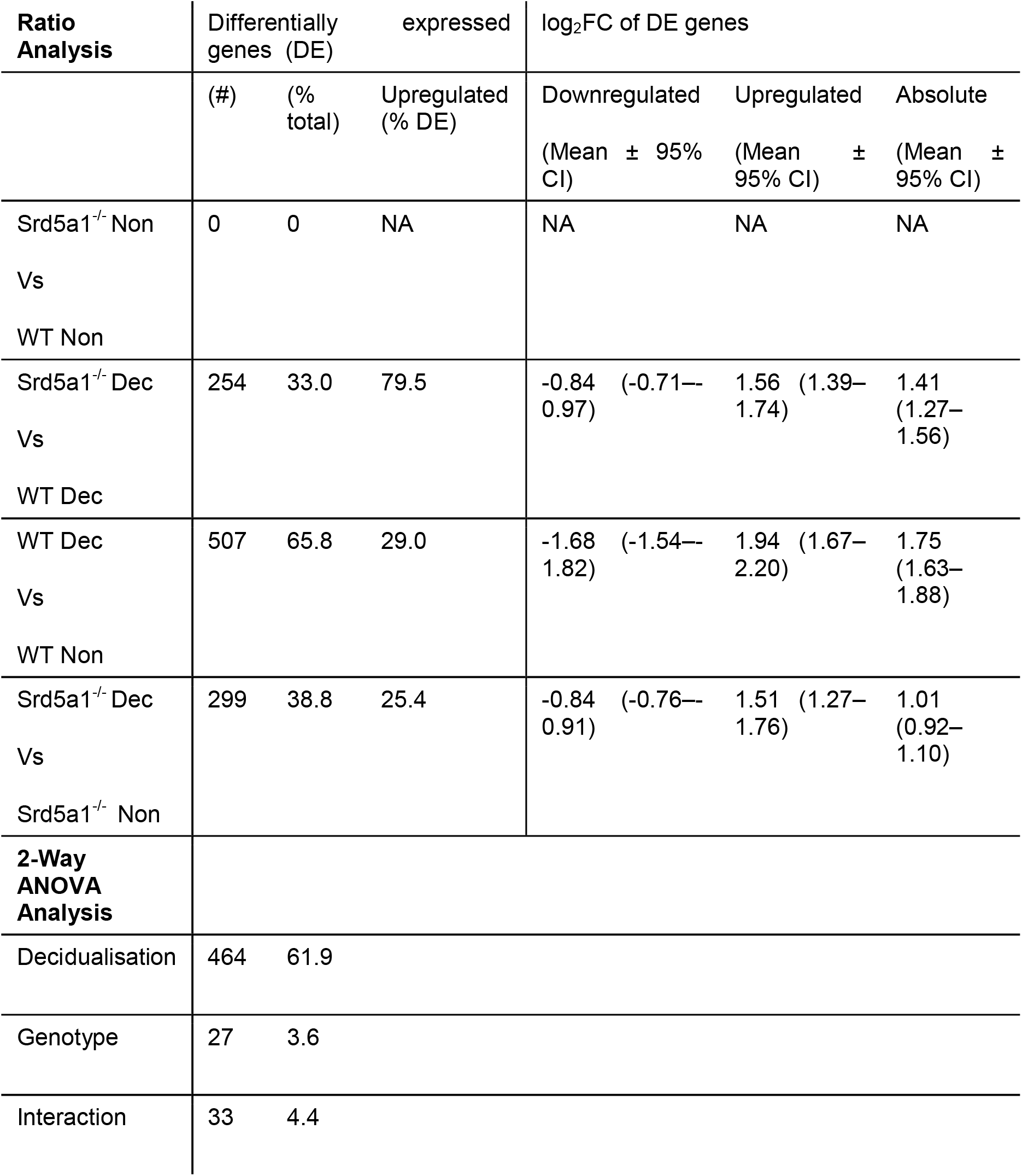
Summary of differentially expressed genes using ratio analysis and 2-Way ANOVA analysis of Pancancer Pathways Nanostring panel. Abbreviations: Non, non-decidualised uterus; Dec, decidualised uterus; DE, differentially expressed; FC, fold-change; CI, confidence interval.

To identify genes differentially regulated by the Srd5a1^-/-^ genotype, we performed two-way ANOVA on Nanostring data using the count number of all genes and the variables decidualisation (Non/Dec) and genotype (WT/Srd5a1^-/-^). Results are shown in Table 1. We identified 47 genes significantly regulated by either genotype or the interaction of genotype and decidualisation, and gene lists can be found in Supplementary Data 2. We further validated key differentially expressed genes from this set by qPCR in a cohort containing additional samples and these tissues showed gene expression patterns that matched those seen in the Nanostring counts or followed similar trends (Fig. 3C).

### In silico analysis of gene networks associated with defective decidualisation in Srd5a1^-/-^ mice identifies deficient expression of angiogenesis regulatory pathways

To gain insight into which gene expression pathways are driving decidualisation, and how they are affected by deletion of *Srd5a1*, we performed gene ontology (GO) analysis based on DE genes identified from our Nanostring data analysis. Upregulated and downregulated genes from WT Dec vs WT non and Srd5a1^-/-^ Dec vs WT Dec comparisons were analysed separately.

No significant GO terms were identified based on genes upregulated by decidualisation (WT Dec vs WT Non), however in the downregulated genes there were several terms related to transmembrane transporter activity (Supplementary Figures 5A). Category netplot (cnet) analysis of the enriched genes showed that all the GO terms were related to the same set of genes (Supplementary Figures 5B). Many of these genes were subunits of the L-type voltage-dependent Ca^2+^ channel (VDCC) which has been shown to be downregulated with decidualisation in endometrial stromal cells (Kusama et al., 2015). In agreement with these findings, many significant GO terms were identified for genes upregulated in Srd5a1^-/-^ decidua compared to WT (Srd5a1^-/-^ Dec Vs WT Dec) all of which related to transmembrane ion transport (Supplementary Figures 5C, D).

**Figure 5.**
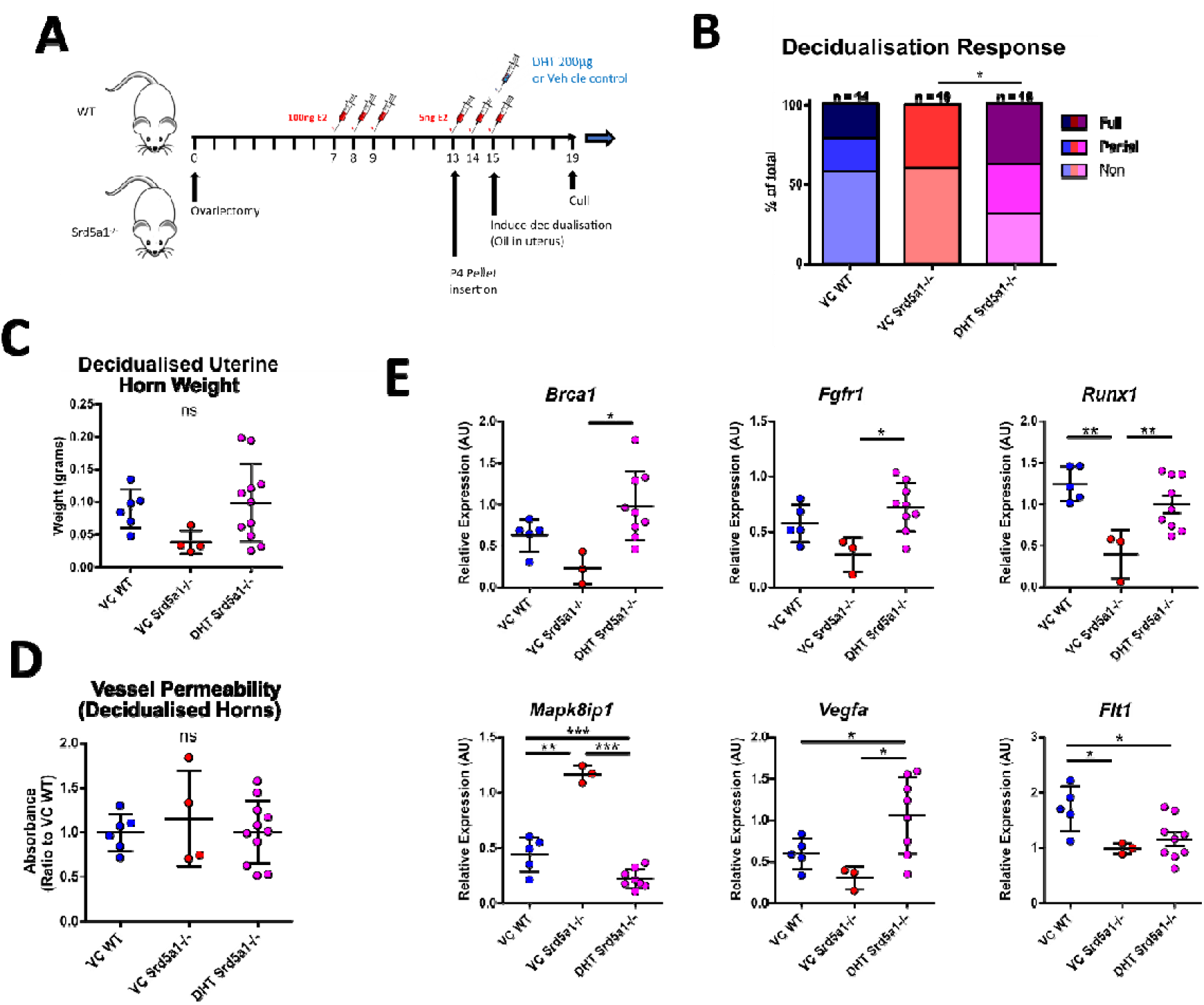
Impact of DHT administration on decidualisation response in Srd5a1^-/-^ mice. A) Schema of decidualisation induction experiment in WT and Srd5a1^-/-^ mice with addition of DHT or vehicle control (VC) at point of decidualisation induction. B) Contingency table depicting proportion of horn responses in WT and Srd5a1^-/-^ mice treated with DHT or VC. C) Wet weight of uterine horns. D) Quantification of Evans Blue dye absorbance in uterine tissue extracts. E) Expression of selected genes analysed by qPCR. Plot in B analysed by chi-squared test; C,D,E analysed by one-way ANOVA and Neuman-Keuls post-hoc tests. * p < 0.05; ** p < 0.01; *** p < 0.001; ns = no significant differences detected.

Significant GO terms were also identified for genes downregulated in Srd5a1^-/-^ decidualised uterus compared to WT (Srd5a1^-/-^ Dec Vs WT Dec ; Fig. 4A). The majority related to cell migration, however there was also a large proportion of GO terms related to the regulation of the vasculature, endothelial cells, or angiogenesis. Cnet analysis highlighted multiple highly connected genes from canonical angiogenesis pathways such as the VEGF pathway (Fig 4B; *Flt1* (VEGFR1), *Kdr* (VEGFR2), *Vegfa, Pgf*). This Cnet also shows that many of the genes related to migration are also linked to angiogenesis pathways. This analysis highlights that gene regulatory pathways that promote angiogenesis are significantly downregulated in decidualised uteri from Srd5a1^-/-^ mice.

### Genes differentially regulated by 5α-reductase deficiency are expressed by stromal and epithelial cell populations in cycling uterus

To identify which cells in the uterus were likely to have altered gene expression as a result of deletion of *Srd5a1*, we compared the results of our Nanostring analysis with our previously published single cell RNA sequencing (scRNAseq) data from cycling mouse uterus which included mesenchymal cells (i.e. fibroblasts and perivascular cells), which are AR^+^ (Simitsidellis et al., 2018, Simitsidellis et al., 2016), and epithelial cells (Kirkwood et al., 2021). Expression of significantly DE genes from the Srd5a1^-/-^ Dec vs WT Dec ratio analysis were sorted by hierarchical clustering (Fig. 4C). All genes had detectable expression in at least one cell population and AR was expressed in stromal (fibroblast>perivascular), but not epithelial, populations. Together, stromal fibroblasts and perivascular cells have detectable expression of the majority of the angiogenic and migratory genes whose expression is regulated by 5α-reductase deficiency identified in the cnet plot (Fig 4B). Perivascular cells play a key role in supporting the growth, stability, and integrity of blood vessels, as well as playing a major role in angiogenesis. This analysis highlights AR^+^ perivascular cells and fibroblasts as plausible cell types mediating the angiogenesis related effects of SRD5A1 activity.

### DHT rescues impaired decidualisation response in Srd5a1^-/-^ mice

Given that AR+ cell types were associated with differential gene expression in Srd5a1^-/-^ mice, we postulated that a lack of intracrine androgen may have altered cell function during decidualisation in the presence of 5α-reductase deficiency. To test whether impaired decidualisation in Srd5a1^-/-^ mice is due to a reduction in intracrine/paracrine DHT, we performed a rescue experiment by administering exogenous DHT coincident with the decidualisation stimulus (Fig. 5A). Following this treatment, the rate of decidualisation in DHT-treated mutants (Srd5a1^-/-^-DHT) significantly increased compared to vehicle-treated mutants (Srd5a1^-/-^-VC) and was statistically indistinguishable from vehicle-treated WT animals (WT-VC; Fig. 5B). Furthermore, the weight of decidualised uterine horns was also the same between WT-VC and Srd5a1^-/-^-DHT groups (Fig. 5C).

We also tested whether the vessel permeability phenotype seen in Srd5a1^-/-^ animals could be rescued by exogenous DHT and found that Evans Blue dye detection in extracts from decidualised Srd5a1^-/-^-DHT was equivalent to that from WT-VC horns, demonstrating that the vascular permeability phenotype is also rescued by DHT administration.

Finally, we tested whether altered gene expression identified in the Srd5a1^-/-^ uterus was affected by administration of DHT. Notably, DHT restored mRNA expression levels to WT-VC levels (*Brca1, Fgfr1, Mapk8ip1, Runx1, Vegfa*; Fig. 5E), and others showed the same trend (*Flt1, Pecam, Il11, Rasgrp1*; Fig. 5E, Supplementary Figures 6A). For some mRNAs there was no evidence of a rescue (*Itga3, Pik3cb, Pik3r2*; Supplementary Figures 6B). The most significant change in gene expression in response to DHT supplementation occurred in genes related to angiogenesis signalling and downstream signal transduction pathways such as *Vegfa, Fgfr1*, and *Mapk8ip1*, suggesting that these stromal/vascular associated pathways are androgen-regulated during decidualisation.

Overall this shows that gross tissue changes and many of the transcriptomic changes associated with impaired decidualisation in Srd5a1^-/-^ mice can be restored by administration of DHT.

## Discussion

Correct and timely decidualisation is essential to the establishment of pregnancy, and defects in decidualisation are linked with implantation failure, pregnancy complications and menstrual disorders such as heavy bleeding (Dunk et al., 2008, Gibson et al., 2018a, Gibson et al., 2016b). We and others have shown that a correct androgen balance is required for optimal decidualisation and fertility (Castracane and Asch, 1995, Cloke et al., 2008, De Vries et al., 1998, Abdalla et al., 1998), but the exact nature of this requirement, e.g. through direct androgen signalling or by acting as precursors to other steroids, and its downstream effects have not been established in an in vivo setting. Identifying the mechanisms through which androgens support the establishment of pregnancy is therefore essential to our understanding of endometrial function and to the development of future fertility treatments. To this end, we have utilised a mouse line and pharmacological approaches to disrupt steroid 5α-reductase activity, which is necessary for the production of DHT, in the endometrium during a model of decidualisation and analysed the effect of this intervention on uterine physiology, transcriptome, and capacity to decidualise.

These investigations have revealed that androgen signalling in the form of locally produced DHT is required for decidualisation to proceed as normal. We have shown that decidualisation is severely impaired in the absence of SRD5A1, the enzyme responsible for local DHT production, and that exogenous DHT administration rescues this effect. Previously, we have shown *in vitro* that hESF upregulate androgen biosynthetic enzymes and produce androgens as they are induced to decidualise, and that AR antagonism blocks intracrine signalling and delays the full differentiation of these cells (Gibson et al., 2016a). The current findings extend this paradigm, demonstrating that intracrine androgen signals are required *in vivo* to drive decidualisation and promote vascular remodelling.

It is possible that lower decidualisation rates observed in Srd5a1^-/-^ mice represent a delay in tissue remodelling compared to WT. In the context of the tightly regulated fertility cycles of mammals such a delay would be sufficient to severely affect fertility. This observation aligns with the reduced fecundity of Srd5a1^-/-^ mice (Mahendroo and Russell, 1999), which may in part be due to decidualisation/implantation defects.

Gene expression changes and the results of the Evans Blue permeability assay suggest a strong role for DHT signalling in vascular remodelling during decidualisation. Decidualised tissue of Srd5a1^-/-^ mice had downregulated expression (vs WT) of gene pathways from two major areas: cell migration and angiogenesis/vascular genes. The process of angiogenesis is inseparable from migration, and migratory genes have previously been shown to be androgen-dependant *in vitro* in endometrial stromal cells (Cloke et al., 2008). It is well established that activated AR promotes, indirectly, VEGF expression and elements of the VEGF pathway (Aslan et al., 2005, Cloke et al., 2008, Joseph et al., 1997, Sordello et al., 1998); and components of this pathway, such as *Flt1* and *Vegfa*, were substantially downregulated in Srd5a1^-/-^ Dec uteri compared to WT. We propose that the vascular phenotypes and impaired decidualisation shown here in Srd5a1^-/-^ mice are a direct result of impaired VEGF and other angiogenic signalling due to a lack of promotion from AR. This paradigm opens up new avenues of exploration for treatments of decidualisation disorders involving impaired androgen signalling as a contributing factor, such as premature ovarian failure, adrenal insufficiency, or in women of advanced maternal age (Vegunta et al., 2020).

In our study finasteride administration did not affect the rate of decidualisation or uterine horn weight in WT mice whereas in Srd5a1^-/-^ mice DHT rescued these features. There are several possible explanations for this discrepancy. The simplest is that, since finasteride was only administered at the point of decidualisation stimulus, sufficient residual DHT was already present in the tissue to support a normal decidualisation response (Rittmaster et al., 1988, von Deutsch et al., 2012). Although finasteride inhibits both type I and II 5α-reductase enzymes in rodents (Livingstone et al., 2015, Thigpen and Russell, 1992) some residual SRD5A1 activity will persist in the presence of finasteride (Upreti et al., 2015) and this may also be sufficient to support decidualisation. Furthermore, its possible that global deletion of *Srd5a1* throughout development and the lifespan of these mutant mice has broader impacts on uterine function that cannot be recapitulated by transient pharmacologic inhibition of the enzyme. Finasteride did however increase vessel permeability during decidualisation in WT mice, similar to the phenotype observed in Srd5a1-/-mice. These findings may represent distinct responses to intracrine androgens such that AR-positive perivascular cells are more sensitive to inhibition of 5α-reductase than stromal fibroblasts during decidualisation and/or that the impact of inhibition on these different cell types is time-dependent. It is notable that endothelial nitric oxide synthase (Nos3), which is a potent, acute mediator of endothelial function, is downregulated during decidualisation in Srd5a1-/-mice (Khorram et al., 1999). This suggests that pathways downstream of intracrine androgens may be acutely or temporally regulated and thus differentially impacted by transient inhibition via finasteride.

A limitation of this study is that targeting the SRD5A1 protein is likely to have effects on steroid hormone signalling beyond its elimination of DHT. In previous work with the same mouse line, the cause of its defective parturition phenotype was identified as a lack of another 5α-reduced androgen, 5α-androstan-13α,17β-diol, indicating there may be specific effects of other 5α-reduced steroids that are not accounted for here (Mahendroo et al., 1999). A separate mechanism by which the Srd5a1^-/-^ mutation may affect results is through diversion of precursor steroids towards other pathways, as occurs during pregnancy when a mid-gestational T surge is aromatised rather than 5α-reduced, causing foetal death due to estrogen excess in Srd5a1^-/-^ mice (Mahendroo et al., 1997). These factors may account for why some gene expression changes were not rescued by DHT administration, and provide an alternative explanation for why finasteride administration does not fully recapitulate the Srd5a1^-/-^ phenotype.

Androgen availability declines with age and there is a concomitant age-related increase in risk of pregnancy disorders (Odibo et al., 2006, Kahveci et al., 2018). The results of the current study highlight the importance of intracrine androgen signalling to early pregnancy remodelling. Given that intracrine androgen action is directly correlated to precursor availability (Gibson et al., 2018b), a decrease or deficiency in androgen precursors could directly impact decidualisation capacity. Further studies are therefore warranted to investigate a possible causal link between androgen deficiency and disorders of pregnancy. Future work in response to this study should investigate intracrine androgen signalling effects in the context of natural pregnancy, for example testing whether DHT can improve fecundity in Srd5a1^-/-^ female mice. It will also be valuable to ascertain the exact mechanism through which AR signalling activates the VEGF pathway, and whether stimulating this pathway can likewise improve fertility in the context of androgen depletion or deficiency. This will facilitate an improved diversity of treatment options for cases where androgen supplementation is not desirable (Vegunta et al., 2020).

In summary, using a well validated mouse model we have confirmed that intracrine androgen signalling, stimulated by locally produced DHT, is required for the robust and timely execution of pathways that stimulate gene expression and changes in cell function essential for a robust decidualisation response. By furthering our understating of the role androgens play in endometrial receptivity and related endometrial disorders these findings will aid in the development of fertility treatments and other interventions.

## Author contributions

I.W.S designed and performed the experiments, analysed the data, interpreted the results, and wrote the manuscript. P.M.K performed the experiments, analysed the data, and interpreted the results. D.R, R.A, F.L.C & L.B.S performed the experiments. D.L provided study materials. P.T.K.S conceived the study, designed the experiments, interpreted the results and revised the manuscript. D.A.G conceived the study, designed and performed the experiments, analysed the data, interpreted the results, and revised the manuscript.

