## Supplementary Figures for "A Role for Steroid 5 alpha-reductase 1 in Vascular Remodelling During Endometrial Decidualisation"

### Slide 1
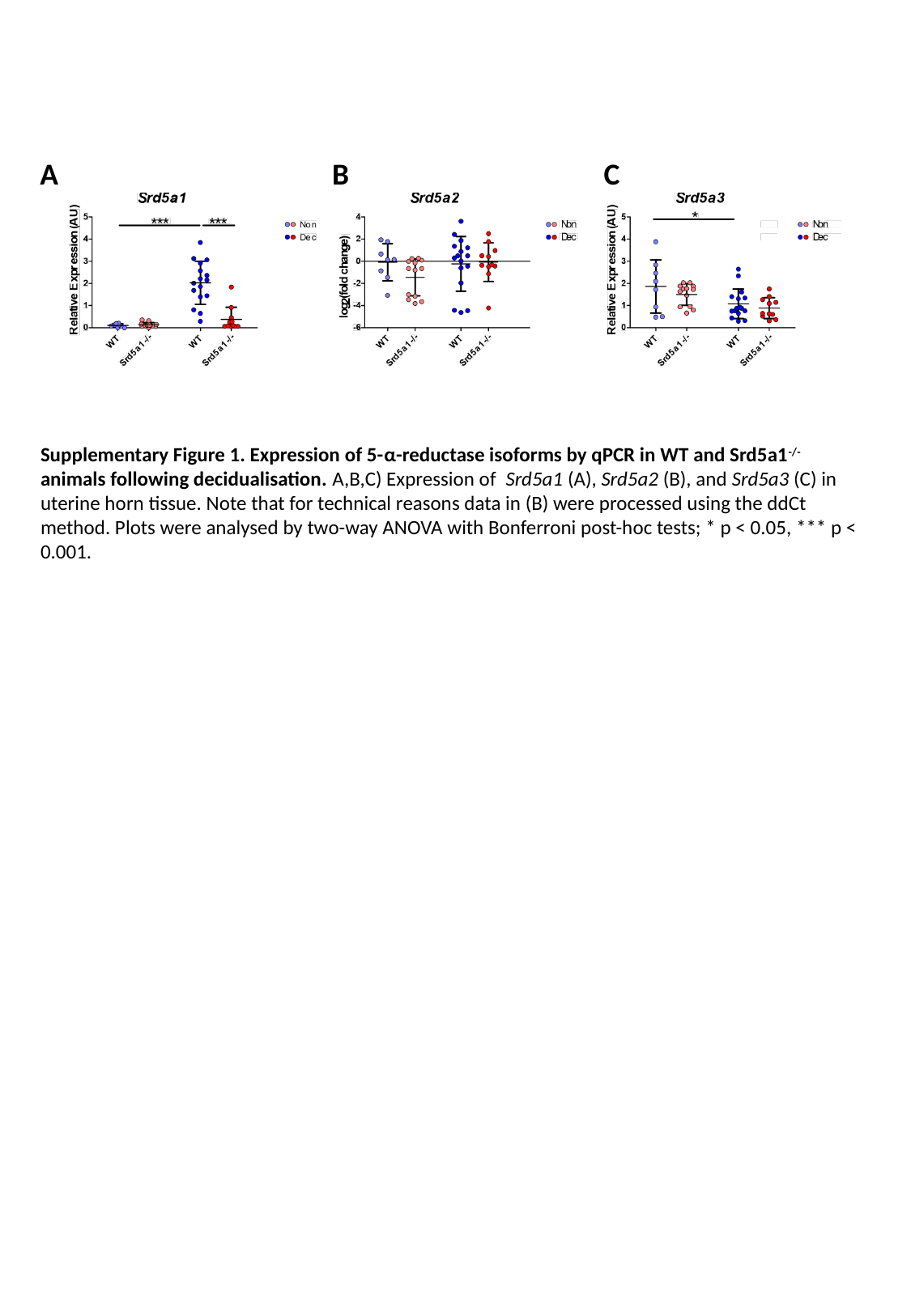

A
B
C
Supplementary Figure 1. Expression of 5-α-reductase isoforms by qPCR in WT and Srd5a1^-/-^ animals following decidualisation. A,B,C) Expression of Srd5a1 (A), Srd5a2 (B), and Srd5a3 (C) in uterine horn tissue. Note that for technical reasons data in (B) were processed using the ddCt method. Plots were analysed by two-way ANOVA with Bonferroni post-hoc tests; * p < 0.05, *** p < 0.001.

### Slide 2
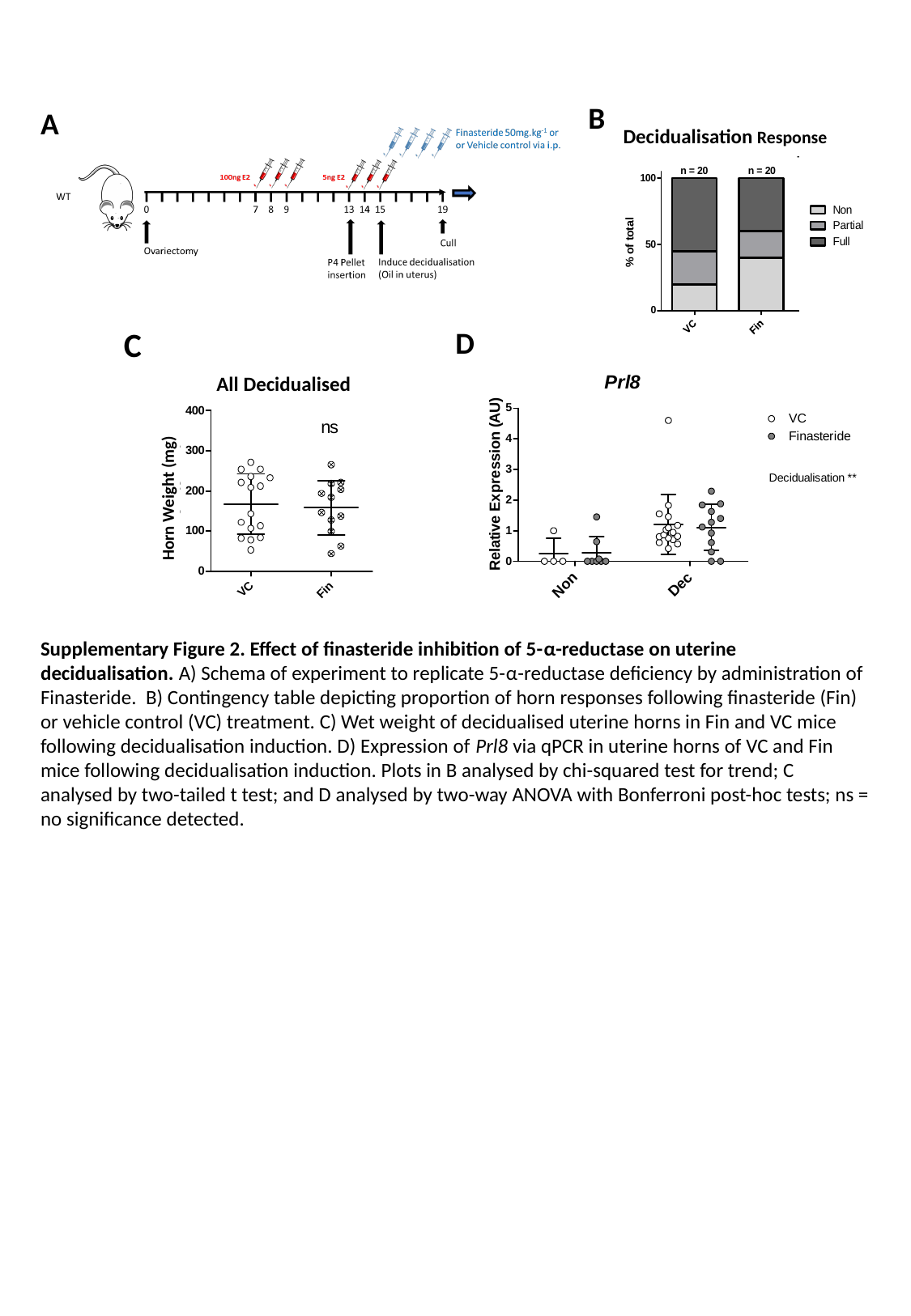

B
Decidualisation Response
A
C
All Decidualised
Horn Weight (mg)
D
Supplementary Figure 2. Effect of finasteride inhibition of 5-α-reductase on uterine decidualisation. A) Schema of experiment to replicate 5-α-reductase deficiency by administration of Finasteride. B) Contingency table depicting proportion of horn responses following finasteride (Fin) or vehicle control (VC) treatment. C) Wet weight of decidualised uterine horns in Fin and VC mice following decidualisation induction. D) Expression of Prl8 via qPCR in uterine horns of VC and Fin mice following decidualisation induction. Plots in B analysed by chi-squared test for trend; C analysed by two-tailed t test; and D analysed by two-way ANOVA with Bonferroni post-hoc tests; ns = no significance detected.

### Slide 3
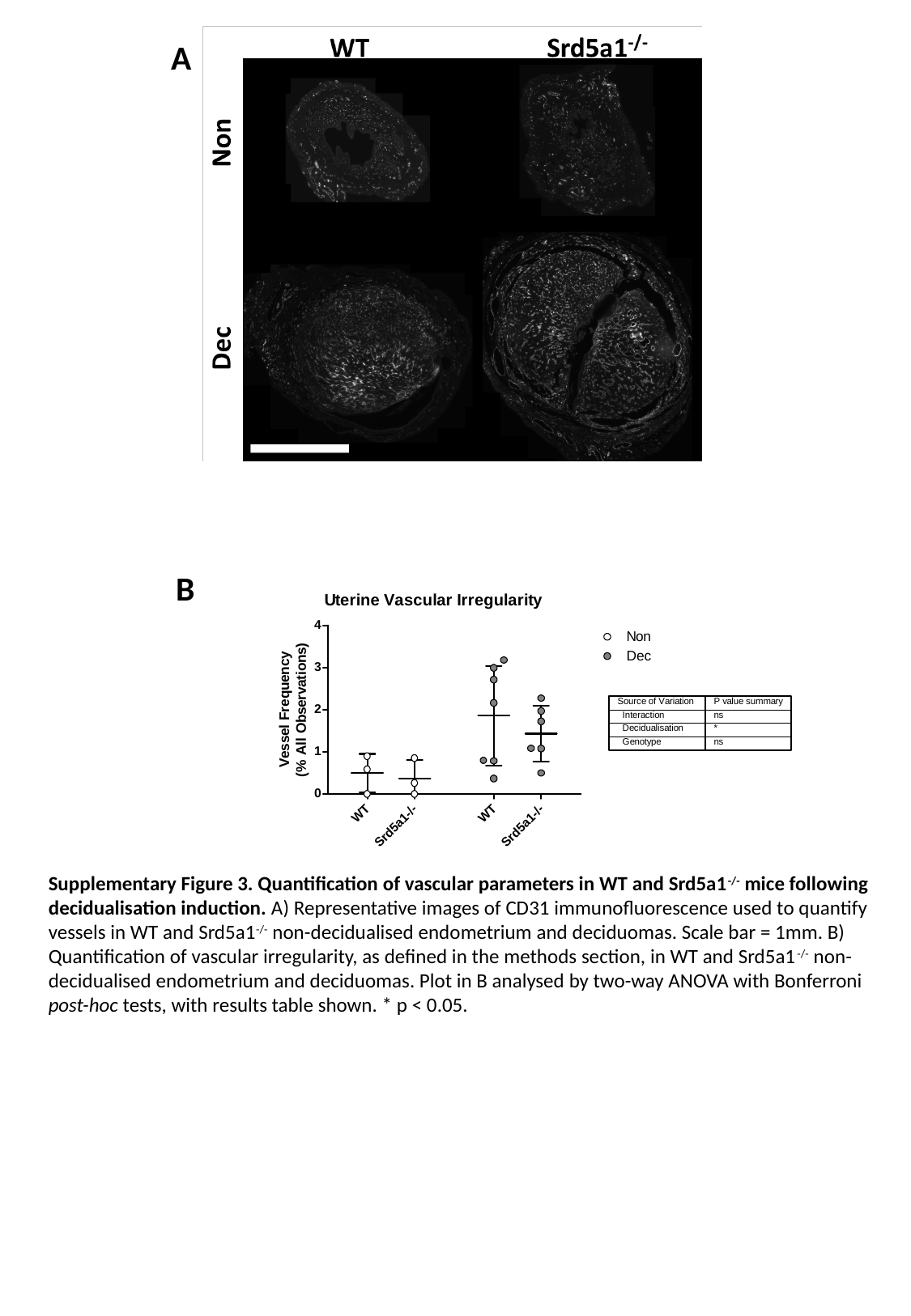

A
Scale bar
B
Supplementary Figure 3. Quantification of vascular parameters in WT and Srd5a1^-/-^ mice following decidualisation induction. A) Representative images of CD31 immunofluorescence used to quantify vessels in WT and Srd5a1^-/-^ non-decidualised endometrium and deciduomas. Scale bar = 1mm. B) Quantification of vascular irregularity, as defined in the methods section, in WT and Srd5a1^-/-^ non-decidualised endometrium and deciduomas. Plot in B analysed by two-way ANOVA with Bonferroni post-hoc tests, with results table shown. * p < 0.05.

### Slide 4
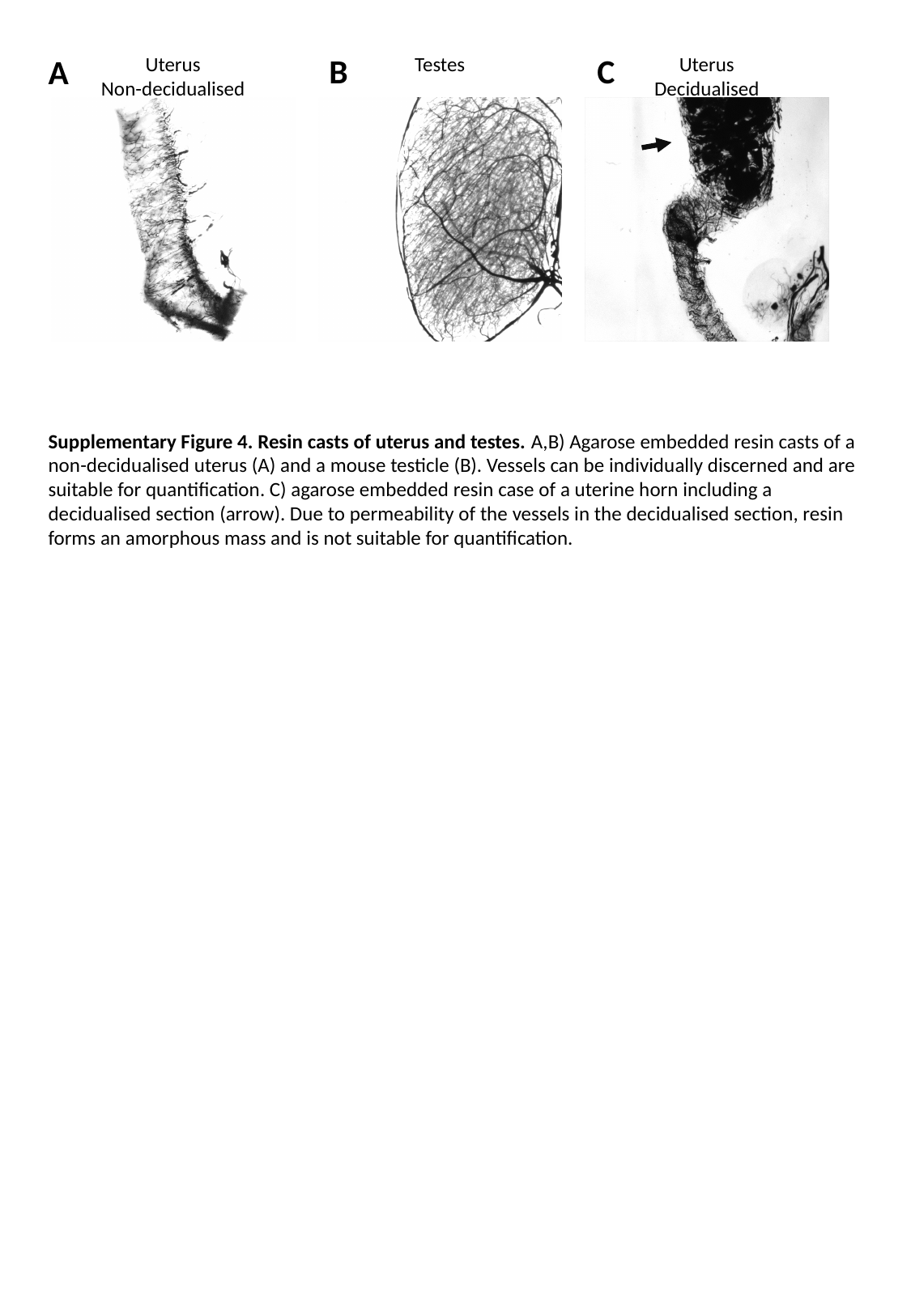

B
C
A
Uterus
Non-decidualised
Testes
Uterus
Decidualised
Supplementary Figure 4. Resin casts of uterus and testes. A,B) Agarose embedded resin casts of a non-decidualised uterus (A) and a mouse testicle (B). Vessels can be individually discerned and are suitable for quantification. C) agarose embedded resin case of a uterine horn including a decidualised section (arrow). Due to permeability of the vessels in the decidualised section, resin forms an amorphous mass and is not suitable for quantification.

### Slide 5
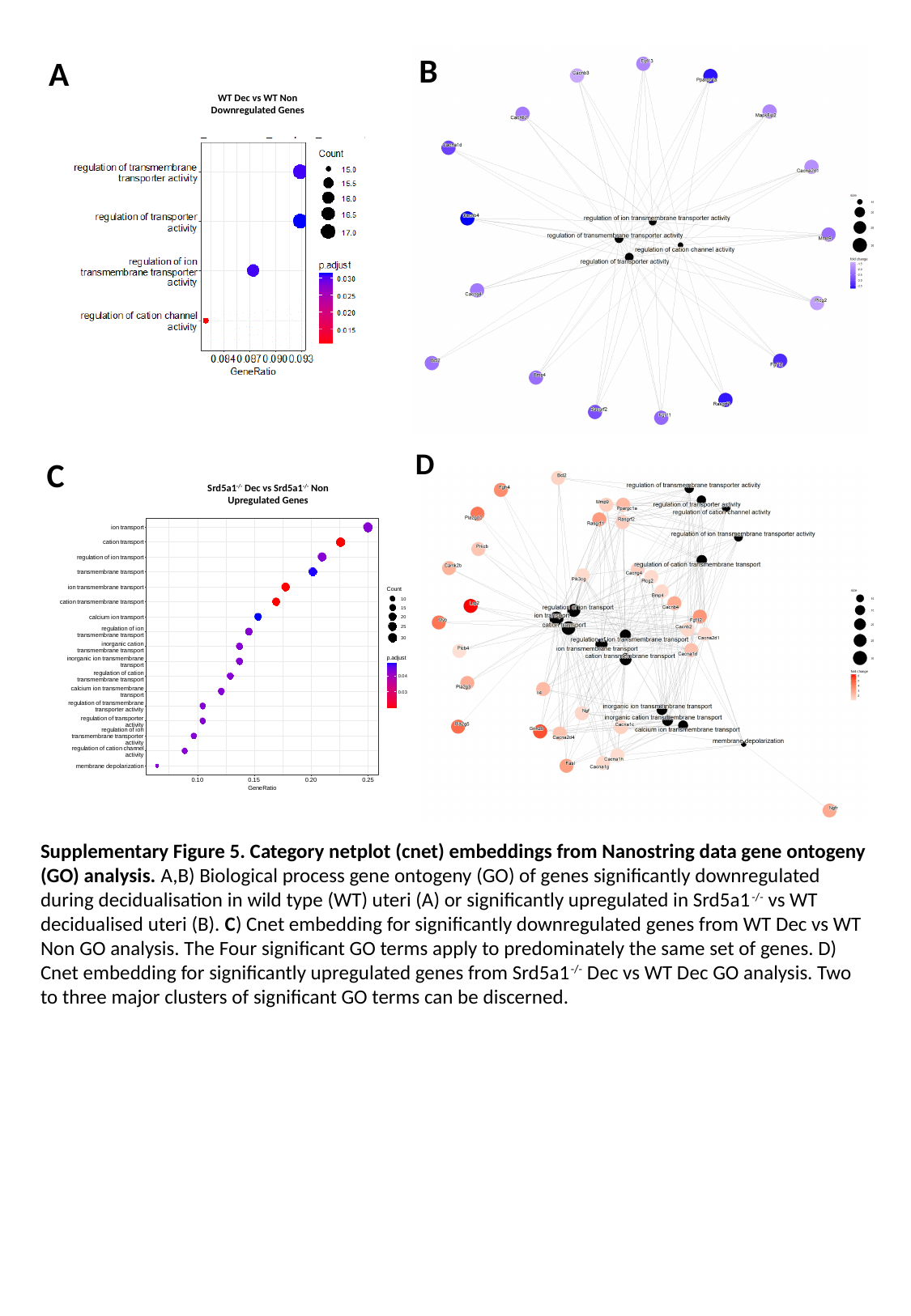

B
C
A
WT Dec vs WT Non
Downregulated Genes
D
C
Srd5a1^-/-^ Dec vs Srd5a1^-/-^ Non
Upregulated Genes
Supplementary Figure 5. Category netplot (cnet) embeddings from Nanostring data gene ontogeny (GO) analysis. A,B) Biological process gene ontogeny (GO) of genes significantly downregulated during decidualisation in wild type (WT) uteri (A) or significantly upregulated in Srd5a1^-/-^ vs WT decidualised uteri (B). C) Cnet embedding for significantly downregulated genes from WT Dec vs WT Non GO analysis. The Four significant GO terms apply to predominately the same set of genes. D) Cnet embedding for significantly upregulated genes from Srd5a1^-/-^ Dec vs WT Dec GO analysis. Two to three major clusters of significant GO terms can be discerned.

### Slide 6
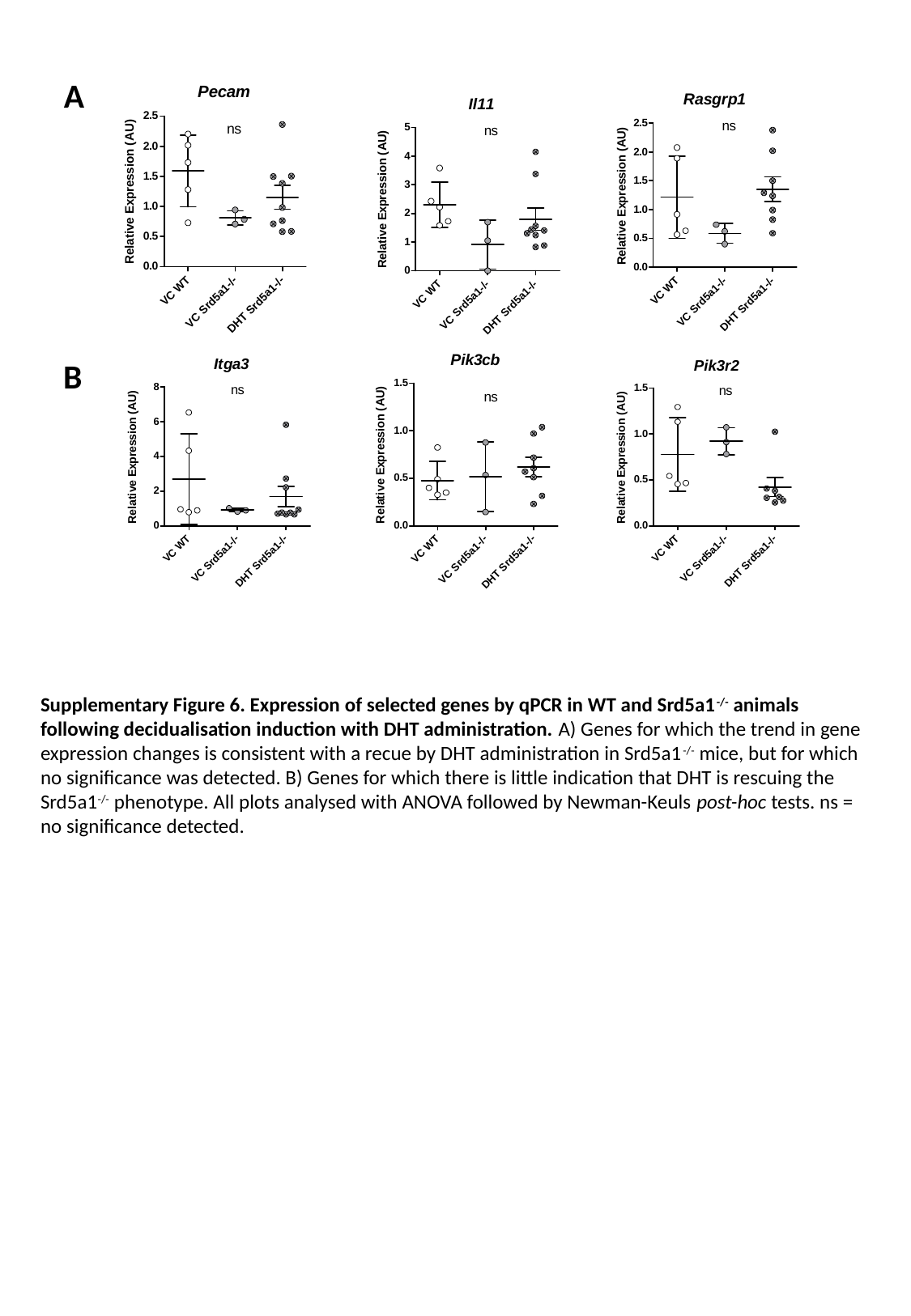

A
B
Supplementary Figure 6. Expression of selected genes by qPCR in WT and Srd5a1^-/-^ animals following decidualisation induction with DHT administration. A) Genes for which the trend in gene expression changes is consistent with a recue by DHT administration in Srd5a1^-/-^ mice, but for which no significance was detected. B) Genes for which there is little indication that DHT is rescuing the Srd5a1^-/-^ phenotype. All plots analysed with ANOVA followed by Newman-Keuls post-hoc tests. ns = no significance detected.
